# Peptidoglycan layer and disruption processes in *Bacillus subtilis* cells visualized using quick-freeze, deep-etch electron microscopy

**DOI:** 10.1101/600171

**Authors:** Isil Tulum, Yuhei O Tahara, Makoto Miyata

## Abstract

Peptidoglycan, which is the main component of the bacterial cell wall, is a heterogeneous polymer of glycan strands crosslinked with short peptides and is synthesized in cooperation with the cell division cycle. Although it plays a critical role in bacterial survival, its architecture is not well understood. Herein, we visualized the architecture of the peptidoglycan surface in *Bacillus subtilis* at the nanometer resolution, using quick-freeze, deep-etch electron microscopy. Filamentous structures were observed on the entire surface of the cell, where filaments about 11-nm wide formed concentric circles on cell poles, filaments about 13-nm wide formed a circumferential mesh-like structure on the cylindrical part, and a “piecrust” structure was observed at the boundary. When growing cells were treated with lysozyme, the entire cell mass migrated to one side and came out from the cell envelope. Fluorescence labeling showed that lysozyme preferentially bound to a cell pole and cell division site, where the peptidoglycan synthesis was not complete. Ruffling of surface structures was observed during electron microscopy. When cells were treated with penicillin, the cell mass came out from a cleft around the cell division site. Outward curvature of the protoplast at the cleft seen using electron microscopy suggested that turgor pressure was applied as the peptidoglycan was not damaged at other positions. When muropeptides were depleted, surface filaments were lost while the rod shape of the cell was maintained. These changes can be explained on the basis of the working points of the chemical structure of peptidoglycan.

## INTRODUCTION

Bacteria, the major inhabitants of the Earth, are in a constant battle to outlast their competitors in the environment and the immune system of host organisms. Most bacterial cells are surrounded by a rigid shield called the “peptidoglycan layer,” which protects them from chemical agents, including lytic enzymes and antibiotics, that are produced by their competitors [1-4].

Peptidoglycan is an essential component of the bacterial cell wall that is found on the outside of the cytoplasmic membrane of almost all bacterial cells except the class *Mollicutes*. This polymer provides strength, rigidity, and shape stability by maintaining turgor pressure. In peptidoglycan, the glycan strands comprise alternating β-1,4-linked *N*-acetylglucosamine (GlcNAc) and *N*-acetylmuramic acid (MurNAc), and the peptide stems are covalently linked to the glycan strands with an amide bond to the carboxyl carbon of the MurNAc. The peptidoglycan layer is also the site of action for antimicrobial agents. Lysozyme, an antimicrobial enzyme critical in animal host defense, is one of the most abundant proteins present on the mucosal surfaces and in body secretions, such as saliva and tears [5, 6]. The epithelial cells secrete lysozyme to protect the host’s mucosal surfaces from infectious bacteria. Lysozyme is also present in white blood cells, especially in granules of phagocytes, where it helps in the elimination of infectious bacteria within phagolysosomes. The underlying mechanism of action of lysozyme involves breaking the bond between GlcNAc and MurNAc (muramidase activity) leading to the degradation of peptidoglycan. On the other hand, fungi and bacteria produce secondary metabolites in defense against predators and competitors [7, 8]. A well-known group of secondary metabolites is β-lactams, a broad class of antibiotics that include penicillin derivatives, cephalosporins, monobactams, and carbapenems. β-lactams inhibit peptidoglycan synthesis by covalently binding to the active site of transpeptidases, known as penicillin-binding proteins (PBPs), cause changes in bacterial cell shape and lead to cell lysis.

This property makes peptidoglycan a vitally important target of β-lactam antibiotics. Therefore, peptidoglycan architecture and the processes that are used to disrupt it are valuable to understand the survival strategies of bacteria and to control pathogenic bacteria. However, the architecture of peptidoglycan is not well understood, because the structure is featured with low density, high flexibility, and is multilayered, which are characteristics that make it unsuitable to be observed using transmission electron microscopy (EM) [2, 9-13].

Quick-freeze, deep-etch replica EM was introduced in order to visualize synaptic transmission processes in 1979, and it has emerged as a useful tool that can be applied for the visualization of many other biological phenomena [14, 15]. It is an advanced technology that is used to visualize biological specimens in an active state as a shot image, with spatial resolution of the order of nanometers and time resolution of sub milliseconds. In this method, the specimen is frozen in less than a millisecond by pressing it against a metal block chilled with liquid helium or liquid nitrogen, resulting in much faster fixation than with chemical methods [15]. Then, the frozen specimen is exposed by fracturing and etching, and shadowed by platinum. The metal shadowing gives a high image contrast, allowing visualization of shot images without image averaging, unlike cryo EM [16, 17]. Moreover, metal contact freezing can be applied to various conditions and specimens, with much more flexibility than with the freezing process for cryo EM. Therefore, quick-freeze, deep-etch replica EM has great advantages, especially when used to visualize low density and flexible structures, in comparison to other methods of transmission EM.

*Bacillus subtilis* is a rod-shaped, Gram-positive, non-pathogenic bacterium that belongs to the phylum *Firmicutes* [1]. The genus *Bacillus* also includes human pathogens such as *Bacillus anthracis* and *Bacillus cereus* [18] and is related to the genus *Clostridium*. Therefore, *B. subtilis* can be an attractive model for the clarification of the architecture and the roles of the cell wall. In this study, the detailed structures of the peptidoglycan layer and its disruption processes in *B. subtilis* were analyzed using the quick-freeze, deep-etch EM and optical microscopy.

## METHODS

### Bacterial strains and media

*B. subtilis* 168 CA and LR2, a strain inducible for L-form derived from *B. subtilis* 168 CA [5, 19] were used. Nutrient agar (NA, Oxoid) was used for routine selection and maintenance of *B. subtilis* 168 CA. Luria-Bertani (LB), nutrient broth (NB, Oxoid), and SMM-defined minimal medium (Spizizen) containing 0.5% xylose or 1 mM isopropyl β-D-1-thiogalactopyranoside (IPTG) were used when required.

### Treatment of cells

Cultured cells around optical density (OD) at 0.2 in 600 nm wavelength, a middle exponential stage, were collected by centrifugation at 8,000 × *g*, 23 °C for 3 minutes. For lysozyme treatment, the cells were suspended in phosphate-buffered saline consisting of 75 mM sodium phosphate (pH 7.3) and 68 mM NaCl to be the original cell density. Lysozyme was added to the cell suspension at final concentration of 0.1 mg/mL and kept at 37 °C without shaking. For Penicillin G treatment, cultured cells were collected and suspended in a fresh medium at the original cell density. PenG was added at the final concentration of 0.1 mg/mL and kept at 37°C with shaking. To induce the L-form transition, 2 × MSM osmoprotective medium (40 mM MgCl_2_, 1 M sucrose and 40 mM maleic acid, pH 7.0) was mixed with the same volume of 2 × NB or 2 × NA. L-form liquid cultivation was done in NB/MSM at 30°C without shaking [19]. Lysozyme hydrochloride, from egg white (Wako Pure Chemical Industries, Osaka, Japan) was labeled with DyLight 488 NHS Ester (Thermo Fisher Scientific, Rockford, IL), according to the instruction.

### Optical microscopy

The cells were inserted into a tunnel chamber with a 5-mm interior width, a 22-mm length, and an 86-μm wall thickness [20]. The tunnel chamber was constructed with a coverslip and a glass slide and assembled with double-sided tape. Fluorescence microscopy was performed with a BX50 fluorescence microscope equipped with a YFP filter unit (U-MYFPHQ) and a phase-contrast setup (Olympus, Tokyo, Japan). Images were captured with a WAT-120NRC charge-coupled-device (CCD) camera (Watec, Yamagata, Japan) and analyzed using ImageJ 1.52.

### Quick-freeze, deep-etch EM

The original cells were collected by centrifugation at 8,000 × *g*, at 23 °C for 5 min and suspended in water (original cells) or a buffer consisting of 10 mM HEPES (pH 7.6), 150 mM NaCl, 1 mM MgCl_2_, and 0.1 mg/mL DNase I (treated cells) to achieve a 20-fold higher cell density. DNase I and MgCl_2_ were included to reduce the viscosity due to bacterial DNA from cells damaged by lysozyme, PenG or *murE* repression. This process was repeated and the cell suspension was mixed with a slurry that included mica flakes, placed on a rabbit lung slab, and frozen by a CryoPress (Valiant Instruments, St. Louis, MO) cooled by liquid helium [21]. The slurry was used to retain an appropriate amount of water before freezing. The specimens were fractured and etched for 15 minutes at −104 °C, in a JFDV freeze-etching device (JEOL Ltd, Akishima, Japan) [22]. The exposed cells were rotary-shadowed by platinum at an angle of 20 degrees to be 2 nm in thickness and backed with carbon. Replicas were floated off on full-strength hydrofluoric acid, rinsed in water, cleaned with a commercial bleach, rinsed in water, and picked up onto copper grids as described [23, 24]. Replica specimens were observed by a JEM-1010 transmission electron microscope (JEOL, Tokyo, Japan) at 80 kV equipped with a FastScan-F214 (T) CCD camera (TVIPS, Gauting, Germany). For tomography, cell replicas were observed by Talos F200C G2 (Thermo Fisher Scientific, Waltham, MA) at 200 kV, and image sets were acquired every degree of angle for 96 steps, by a complementary metal-oxide-semiconductor (CMOS) camera (Ceta camera, FEI) [25]. The images were analyzed by ImageJ.

## RESULTS

### Surface structure of *Bacillus subtilis*

To visualize the structure of the peptidoglycan layer, *B. subtilis* was observed for the first time by quick-freeze, deep-etch EM. Cells in the exponential growth phase were collected by centrifugation and placed on glass, frozen, fractured, deeply etched, shadowed with platinum, and then the platinum replicas were recovered and observed. The shape and dimensions of the cells were consistent with the images of living cells obtained by optical microscopy (Fig. 1A, B). Filamentous structures were clearly observed on the cell surface and can be distinguished between the cell pole and the cylindrical part of the rod-shaped cells (Fig. 1C, D and Movie S1). Thick filaments were aligned in a partial circumferential manner on the cylindrical part and thin filaments were concentrically arranged around both poles. The filament widths were measured based on the image profile, assuming that the mid density of the image profile positions at the boundary of the filament being focused on (Fig. S1). As the structures were coated by platinum to be 2 nm in thickness, the original filament width should be 4 nm thinner than the value obtained. Thus, we concluded that the widths of thick and thin filaments were 8.7 ± 0.4 and 6.9 ± 0.3 nm, respectively, and the thin filaments were aligned with a pitch of 11.3 ± 0.4 nm (Fig. 1E, F).

**Fig. 1.**
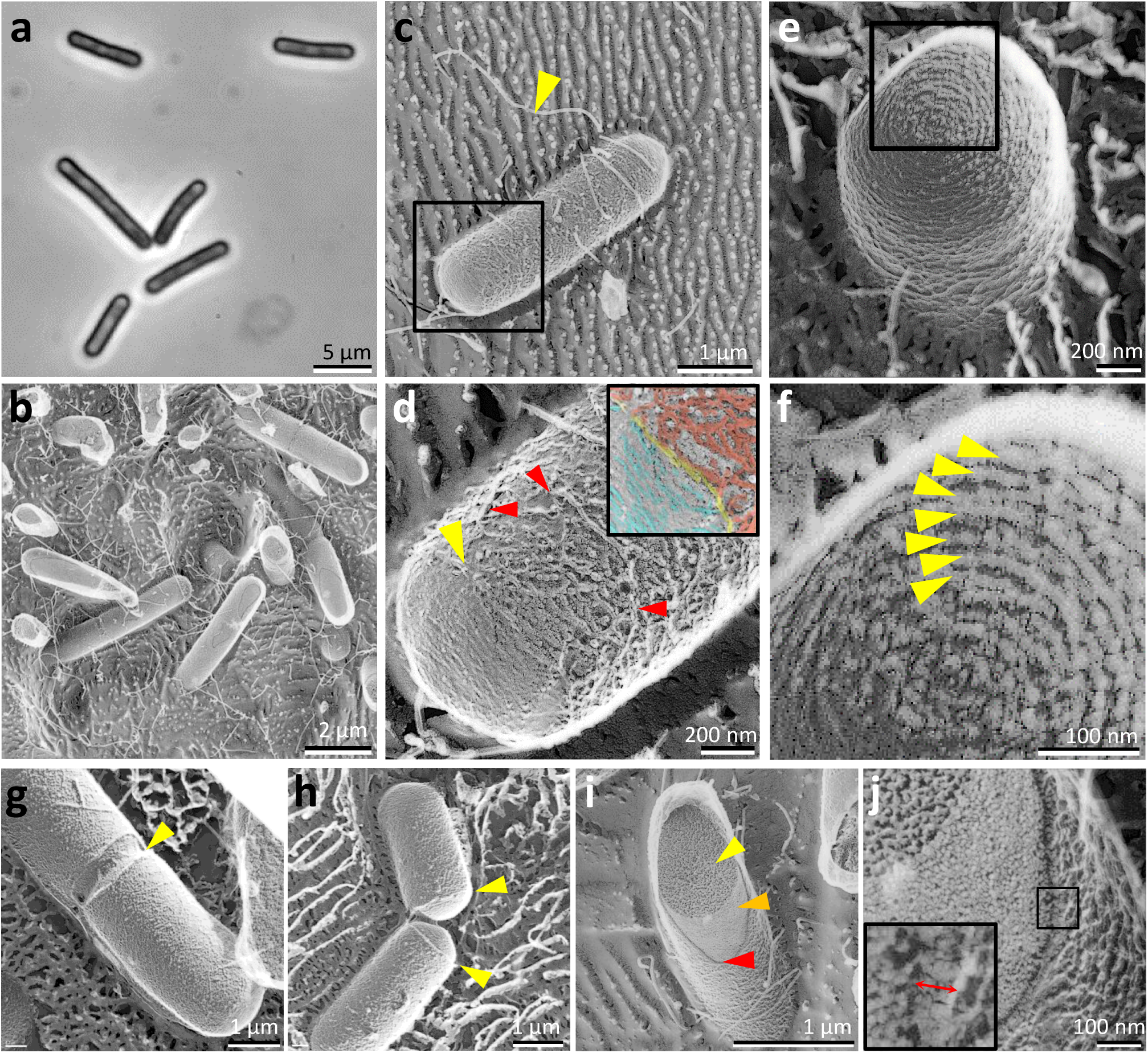
Surface structures of *Bacillus subtilis* 168 CA cells visualized by quick-freeze, deep-etch EM. **A)** Phase-contrast optical microscopy of cells. **B)** Field image of cells. **C-H)** Magnified cell images. **C**) Side view. A flagellum is marked by a blue triangle. A boxed area is magnified in (D). **D)** Magnified cell surface image. Filaments on the cylindrical part are marked by red triangles. The boundary between the cylindrical and pole parts is marked by a yellow triangle. The area positioning at the yellow triangle is shown in an inset and colored for the filamentous structures at cell pole (blue), piecrust (yellow), and cylindrical part. **E**) Pole view. A boxed area is magnified in (F). **F)** Magnified pole surface image. Concentric circles are marked by yellow triangles. **G)** Dividing cell. The invagination is marked by a green triangle. **H)** Daughter cells, probably just after cell division. Piecrust structures are marked by red triangles. **I)** A fractured cell showing three different surfaces. The cytosol, cell membrane, and a cross section of the peptidoglycan layer are marked by yellow, orange, and red arrows, respectively. **J)** Magnified image of fractured part in (I). A cross section of the peptidoglycan layer is marked by an arrow.

On elongated cells prior to cell division, invagination was observed at the central position of the axis, and an obvious boundary was found between the cylindrical part and the surface of the invagination (Fig. 1G). Obvious filamentous structures were not found on the surface of the invagination. A wall 24.8 ± 1.6 nm wide was observed between the surface of the invagination and the cylindrical part (Fig. 1G). This small wall was also found in cells after division (Fig. 1H), but not in isolated single cells (Fig. 1D).

Cells aligned vertically were frequently fractured by a knife, and a fractured cytosol, membrane surface, and cross section of the peptidoglycan layer were visualized (Fig. 1I, J). The height of the peptidoglycan layer was estimated as 30.5 ± 4.1 nm (n = 26).

### Peptidoglycan disruption by lysozyme

Next, we examined how the peptidoglycan layer is disrupted by lysozyme. We added lysozyme from the egg white to growing cells, and observed the changes occurring on the cell structures by phase-contrast optical microscopy (Fig. 2A). The effects of lysozyme on *B. subtilis* cells were consistent with previous reports, although we traced the changes in more detail [5, 26, 27]. In the present observation, the cell mass started to detach from the envelope structure around 30 minutes after the addition of lysozyme and moved to one side over time. Finally, all the mass was localized as a sphere at one side of the cell envelope. After a 60 minute treatment, 100% of the cells (n = 72) were converted to protoplasts.

**Fig. 2.**
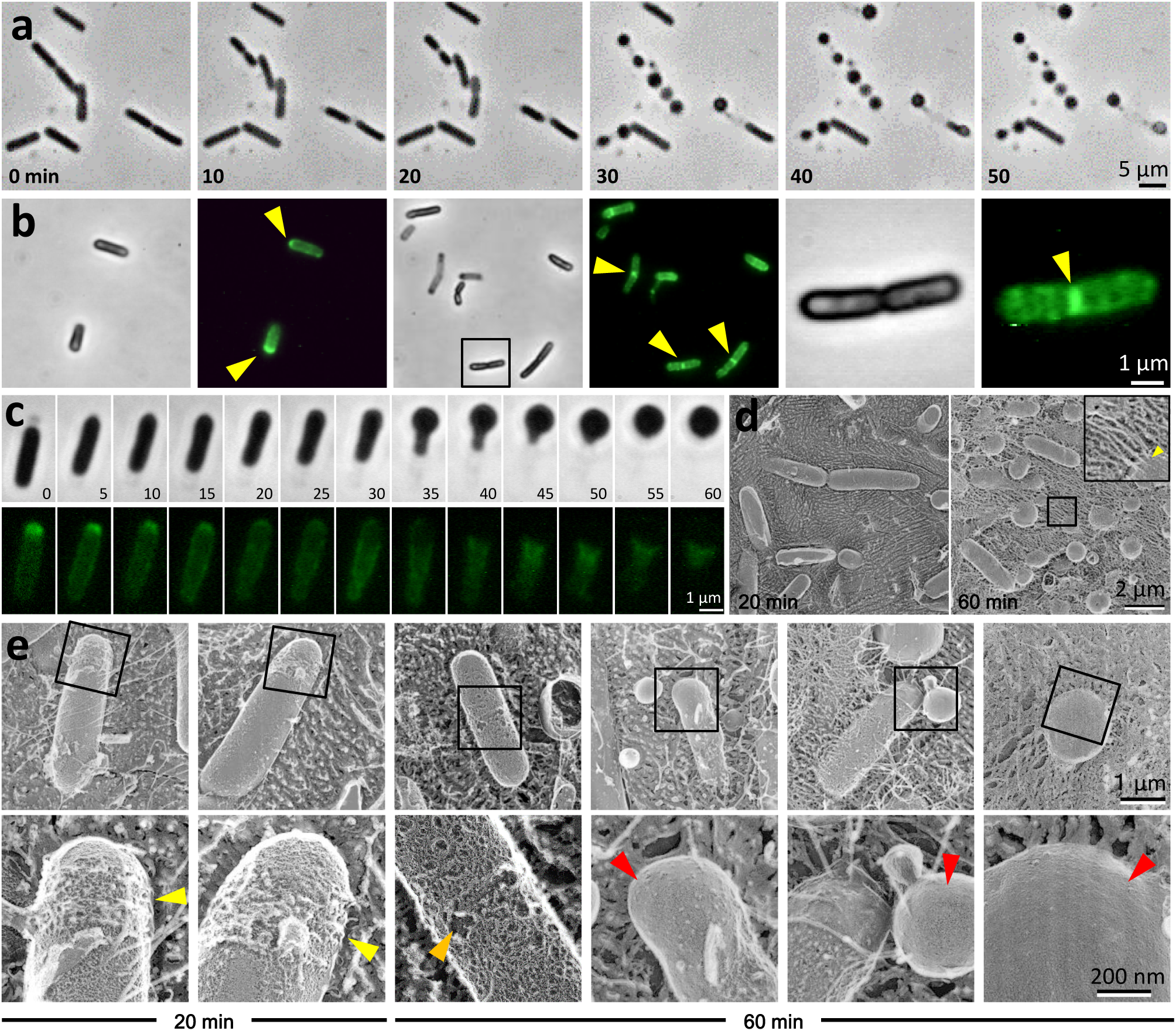
Damages on cell structure by lysozyme. **A)** Phase-contrast microscopic images after addition of lysozyme taken with 10 minutes intervals. **B)** Localization of fluorescently labeled lysozyme on cell. Fluorescence of labeled lysozyme was visualized at 10 minutes after the addition. Phase-contrast and fluorescence images are shown as the left and the right panels, respectively, in each of three image sets. A boxed cell in the middle-paired panels is magnified in the right panel set. The signals were found preferentially at a pole and the division site as marked by yellow triangles. **C)** Phase-contrast and fluorescence images of a single cell treated by labeled lysozyme taken at 5-minute intervals. **D)** Field image of cells treated with lysozyme for 20 and 60 minutes taken by the quick-freeze, deep-etch EM. The mica surface is covered by eutectics appearing as thin filaments. A typical eutectic is shown in the inset with an arrow. **E)** Quick-freeze, deep-etch EM images of cells treated with lysozyme for 20 and 60 minutes, as shown in the bottom. The focused regions boxed in the upper panels are magnified in the lower panels. Ruffling, coarse pattern and smooth surfaces are marked by yellow, orange and red triangles, respectively.

To locate the sites where lysozyme works on the cell in this process, we labeled lysozyme fluorescently with DyLight 488 through amino groups of protein and using fluorescence microscopy, traced its attachment point (Fig. 2B). After 10 minutes, the labeled lysozyme bound to the whole cell surface but preferentially bound to the division site and a cell pole. The changes in lysozyme localization over time were traced through live imaging (Fig. 2C). The signal found at a pole decreased with time and disappeared at 40 minutes. The signal on parts other than the cell poles remained for 40 minutes, and then it moved to one side. The disappearance of lysozyme signal should suggest the dissociation of lysozyme from envelope, caused by the complete digestion of the target sites on the peptidoglycan layer.

Next, we observed the structural changes on the cell surface at high resolution using the quick-freeze, deep-etch EM (Fig. 2D, E). The images featured filamentous structures covering the entire field. They are likely to be filaments of eutectic mixtures, resulting from the cytosol released from damaged cells, because generally, some ions and polymers form filaments of eutectic mixtures in the quick-freeze, deep-etch EM [28]. This assumption is consistent with the fact that the covering of filaments increased with the time of treatment with lysozyme, and PenG and *murE* repression (see below). Eutectic filaments can be confused with the real surface in images; however, they can be distinguished because eutectics are continuous from a position distant from the cell. At 20 minutes after the addition of lysozyme, ruffling of the surface structure was observed around a cell pole, which is probably corresponding to the area to which lysozyme bound on the cell. The peptidoglycan layer should detach from the cell partly by the degradation of the sugar chain. At 60 minutes, the ruffling parts became restrictive, which is consistent with the observation that the area of lysozyme binding became restricted in the fluorescence microscopy. Instead of ruffling, we found coarse patterns on the cylindrical part of cells and spherical structures with a smooth surface, suggesting that lysozyme moved and digested the peptidoglycan at the cylindrical parts.

### Peptidoglycan disruption by penicillin

The process of peptidoglycan disruption by the effect of penicillin was examined (Fig. 3). Beginning 30 minutes after the addition of 100 µg/mL Penicillin G (PenG) to the growing culture, a less dense, flexible structure was observed to project from a side position on a cell under optical microscopy (Fig. 3A). At 120 minutes, 89% of the cells (n = 172) were released from the peripheral structure probably as a protoplast.

**Fig. 3.**
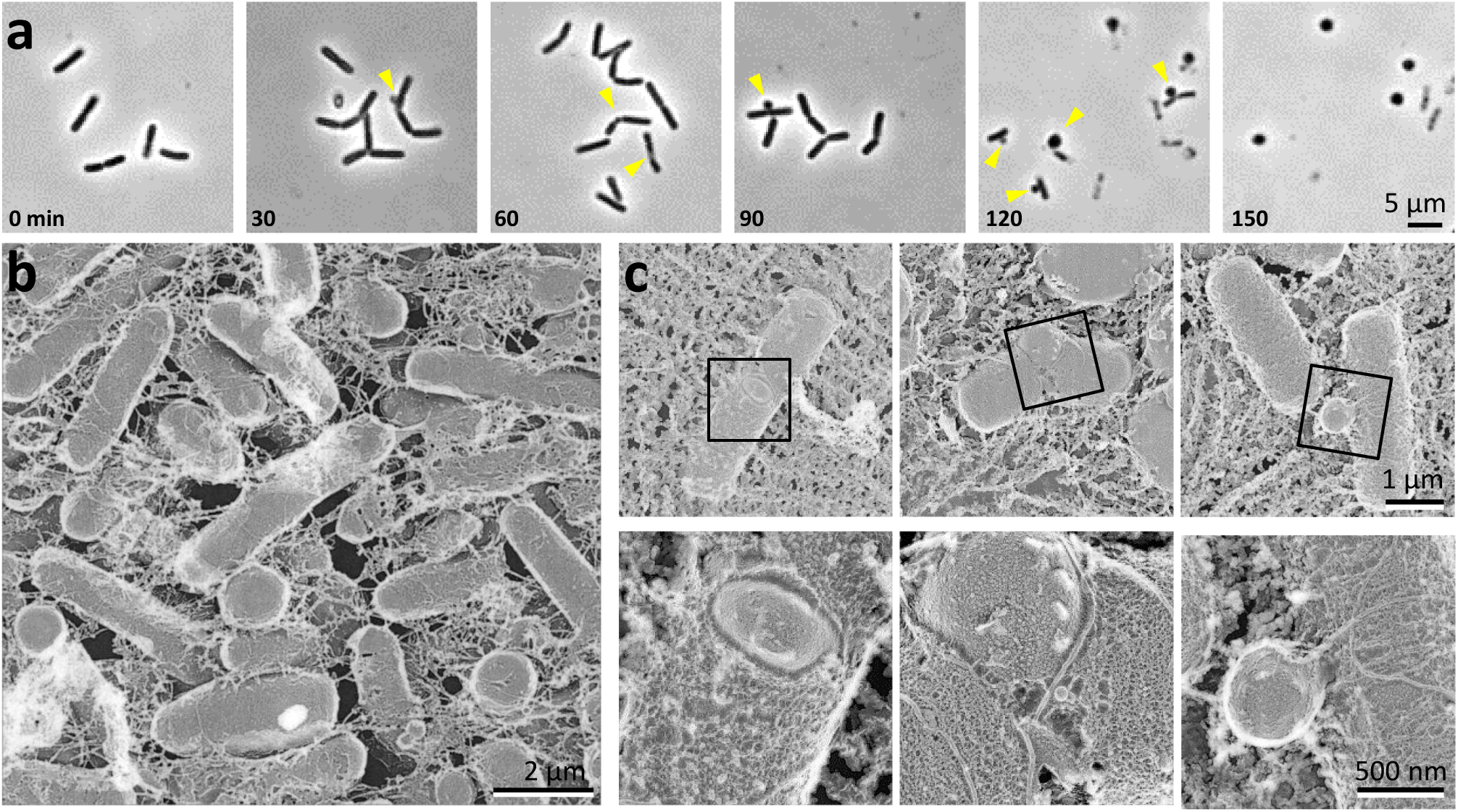
Damages on cell structure by PenG. **A)** Phase-contrast microscopic images after addition of PenG taken at 30-minute intervals. **B, C)** Quick-freeze, deep-etch EM images of cells treated with PenG for 120 minutes. **B**) Field image. **C**) Magnified cell images. The boxed regions in the upper panels are magnified more in the lower panels.

Next, the cells damaged by PenG treatment for 120 minutes were observed by the quick-freeze, deep-etch EM (Fig. 3B, C). The cell images showed various stages of the process by which a protoplast comes out from the cell envelope. In many cases, the cytoplasm came out from the central position of the cell where the cell division should be scheduled (Fig. 3C). Unlike the effect of lysozyme, the cytoplasm in the early stage of cell disruption seemed to be subjected to strong turgor, because the protoplasts coming through the cleft had some outward curvature.

### Peptidoglycan disruption by *murE* operon repression

The repression of the *murE* operon ceases supplying muropeptides, induces the gentle degradation of the peptidoglycan layer, and then results in the transfer of *B. subtilis* cells into L-form [29, 30]. This process was visualized by phase-contrast optical microscopy (Fig. 4A). When the *murE* operon was repressed for 80 minutes, the cell morphology was disturbed mostly around cell poles, featured by a tapered pole, and the detachment of cell mass from the pole. Time lapse imaging of cells showed that these two features occurred in a sequential manner. In those cells, finally the cell became spherical, which may proliferate as L-form. These processes agree with those previously reported [29, 30]. Next, we visualized the cells by the quick-freeze, deep-etch EM. Irregular cell shapes were obvious under this method, including asymmetric cell division, tapered cell pole (Fig. 4B), and spherical extension at a cell pole (Fig. 4C, left). The filaments on the cell surfaces were unclear compared to the original cells, or totally lacking (Fig. 4C, left and middle). On the spherical cells, no filaments were found on the surface, supporting our assumption that the filaments observed in the quick-freeze, deep-etch EM are derived from the peptidoglycan layer (Fig. 4C, right).

**Fig. 4.**
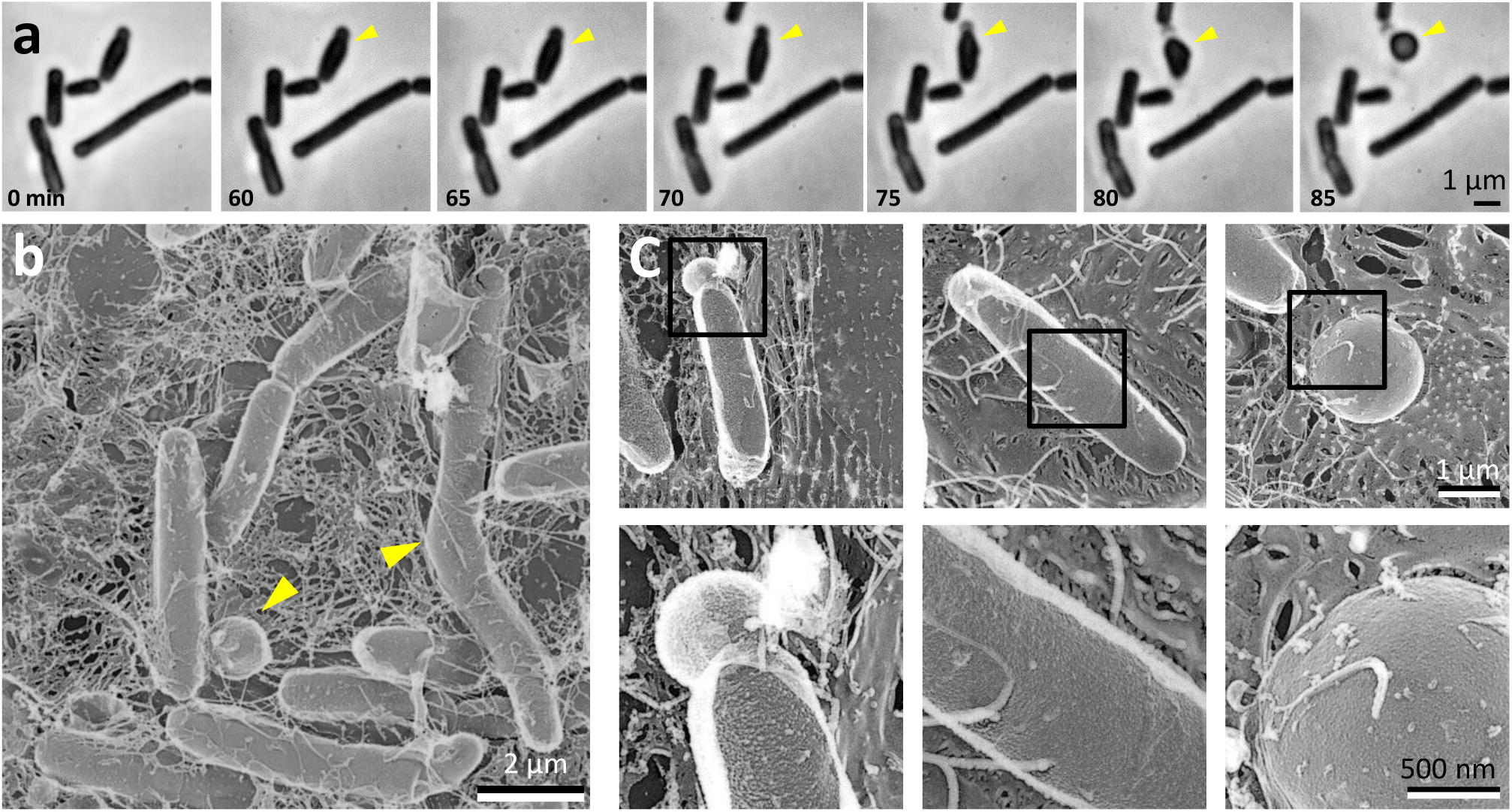
Effects on cell structure by *murE* operon repression. **A)** Phase-contrast microscopic images taken at 5-minute intervals commencing at 60 minutes after repression. The cell marked by a yellow triangle changed its structure drastically. A tapered polar, a bulge, and swollen cells can be observed sequentially with time on the cell. **B, C)** Quick-freeze, deep-etch EM images of cells repressed for *murE* for 85 minutes. **C**) Cell features resulted by *murE* repression. The area boxed in the upper panels are magnified in the lower panels.

## DISCUSSION

### Visualization of peptidoglycan layer by quick-freeze, deep-etch EM

The images obtained by the quick-freeze, deep-etch EM may appear like those from Scanning Electron Microscopy (SEM), but they are featured much higher spatial resolution and sub-millisecond time resolution [14, 15]. This method should be very efficient for research of microorganisms whose interaction with the environment on their surface is critical for many activities [15, 31, 32]. In the present study, the quick-freeze, deep-etch EM was applied to the visualization of the peptidoglycan and its disruption process, for the first time. The results showed the details of filamentous structures at nanometer resolution and their change, as expected, with time resolution. In previous studies, peptidoglycan layers have been visualized by other methods including negative-staining EM, cryo electron tomography, sectioning EM, SEM, AFM, and freeze fracture EM. However, the quick-freeze, deep-etch EM has advantages over other methods. As negative-staining EM and cryo electron tomography need to isolate the peptidoglycan layer, the alignment of the structure on the cell cannot be observed [10, 33]. Sectioning EM does not show the images three dimensionally [34, 35]. SEM can provide three dimensional images but chemical fixation may modify the original structure and the resolution of SEM is much lower than that of the quick-freeze, deep-etch EM [14, 15]. Atomic force microscopy (AFM) visualizes the surface three dimensionally but cross-section images are not available and the resolution is lower than that of the quick-freeze, deep-etch EM, especially when the target structure is vertically aligned against the surface [12, 36]. Although the image obtained with freeze fracture EM is similar to that of the quick-freeze, deep-etch EM, the chemical fixation involved in the former may modify the original structure [37].

In normally-growing cells, the alignments of the peptidoglycan filaments were clearly distinguished between the poles and the cylindrical part (Figs. 1 and 5A left). This was consistent with the knowledge from molecular biology that peptidoglycan synthase is localized by MreB when cells are elongating and the synthase is localized at the division site by FtsZ during cell division [38, 39]. The concentric pattern in the pole is considered to be common to the structure previously observed by a related method, the freeze fracture of fixed cells of *Staphylococcus* [37]. Atomic Force Microscopy (AFM) of the peptidoglycan showed circular pattern for *B. subtilis* [12] and also for *Lactococcus lactis* [11]. The circular pattern observed in the present study may be general in *Firmicute* bacterial species, although the appearance depends on the visualizing methods. A small wall structure was seen at the boundary between the surfaces of the invaginating part and the cylindrical part (Fig. 1G, H). This structure should correspond to a structure named “wall band” observed in sectioned EM images [34, 35]. A similar structure is known as “piecrust” in SEM observation of *Staphylococcus aureus* [37], although it has not been observed in SEM images of *B. subtilis* [2]. In previous observations of *B. subtilis* using SEM, the low resolution and chemical fixation processes interfered with the visualization of the piecrusts. The filament pattern observed on the cylindrical part is a circumferential mesh-like structure similar to the pattern observed for the isolated peptidoglycan layer of *Escherichia coli* using AFM, although the filament widths are different [9]. Perhaps, the filaments in the cylindrical part may be aligned roughly in a circumferential way, generally in rod shaped bacteria [10, 12, 13].

**Fig. 5.**
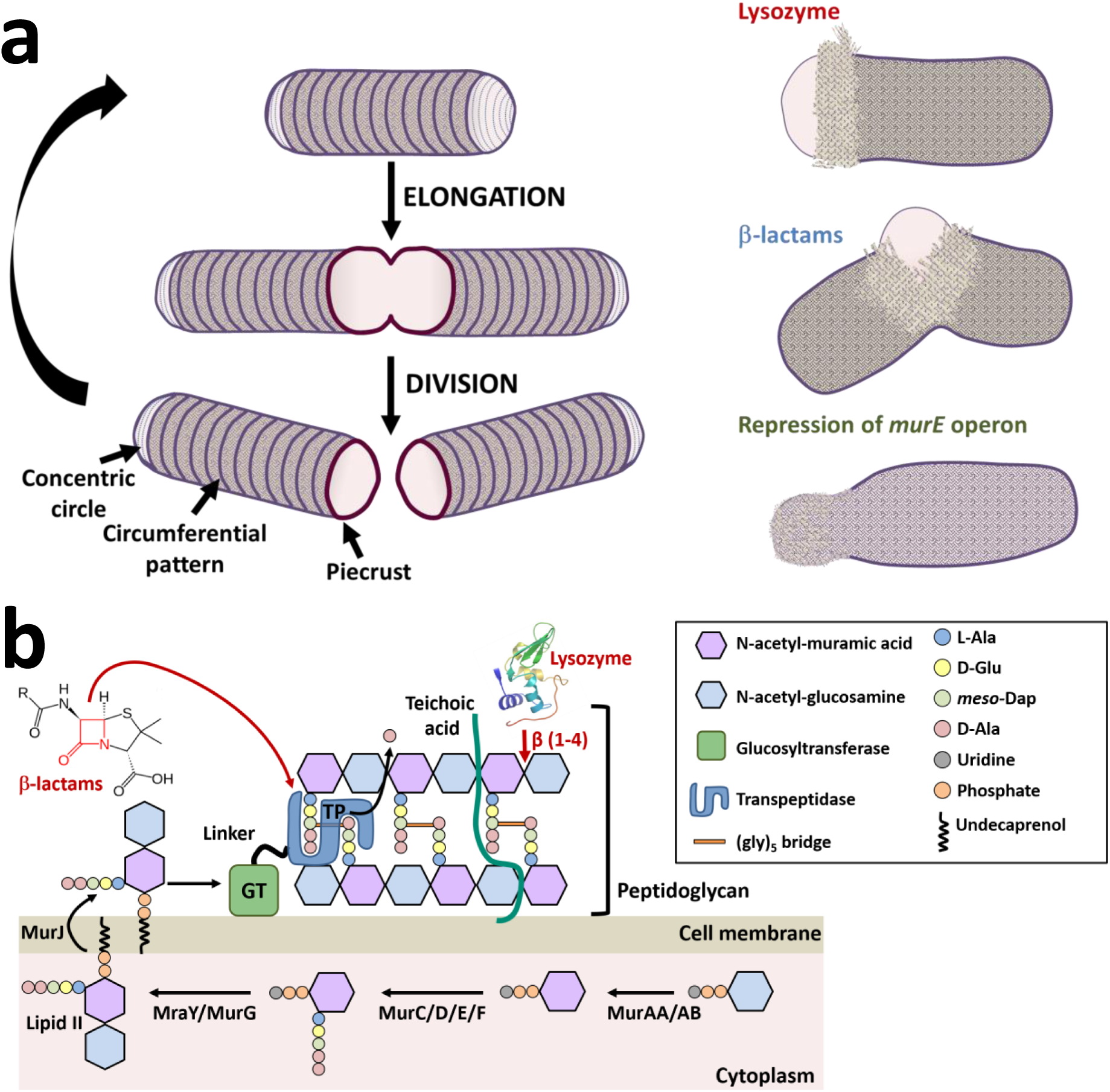
Schematic presentations of peptidoglycan architecture and working points of inhibitory factors. **A)** Structural features on surfaces of cells in the division cycle (left) and peptidoglycan disruption (right). **B)** Chemical processes. The peptidoglycan layer consists of strands of repeating GlcNAc and *N*-acetylmuramic acid (MurNAc) subunits. Transpeptidase (TP) forms a crosslink between two sugar chains, forming peptidoglycan network. Glycosyltransferase forms linkages between GlcNAc and MurNAc residues. Peptidoglycan synthesis starts in the cytoplasm, where the nucleotide precursors are synthesized by the Mur enzymes (MurA, MurB, MurC, MurD, MurE and MurF). MurNAc-pentapeptide is coupled to a membrane carrier, undecaprenyl pyrophosphate, by MraY. Then, lipid II is formed through coupling with GlcNAc by MurG, and transferred to the outside of the cytoplasmic membrane. The antibiotic β-lactams represented by PenG bind to and inhibit the activity of the transpeptidase by forming a highly stable penicilloyl-enzyme intermediate. Lysozyme cleaves the bond between MurNAc and the fourth carbon atom of GlcNAc. The depletion of MurE blocks the supply of muropeptides.

Based on the observation here, we can suggest a scheme for cell surface structures in cell division cycles (Fig. 5A left).

### Lysozyme starts from new pole and division site

In the present study, the disruption process of the peptidoglycan layer was visualized for each of the three factors with different working points (Fig. 5A right, B). In the process, lysozyme preferentially bound to a cell pole and a cell division site (Fig. 2B). Probably, the newly synthesized peptidoglycan has many gaps, which lysozyme molecules can access easily. The tracking of lysozyme showed that it detached from the initially bound position earlier than it did the other parts, suggesting that the peptidoglycan digestion was completed at the cell pole (Fig. 2C). Even when pathogenic or parasitic bacteria invade the host, the division site and the new pole is preferentially attacked by lysozyme. We focused on *B. subtilis* inhabiting the environment in the present study, but the process by which lysozyme attacks pathogenic or parasitic bacteria should also be very interesting.

Quick-freeze, deep-etch EM showed that the action of lysozyme loosens the peptidoglycan. As lysozyme cleaves carbohydrate chains that mainly form the filaments of the peptidoglycan layer (Fig. 5B) [5, 6], the peptidoglycan layer separates from the cell surface (Fig. 2E). This damage on the entire peptidoglycan layer is as effective as the protection system that is involved in the resistance against invasion by pathogenic or parasitic bacteria.

### Turgor by penicillin treatment

In the early stages of cell disruption by PenG, large turgor appeared to be applied to the cytoplasm at the cell division site from the cell inside, because the cell membrane at a cleft of the cell envelope showed curvature to outside (Figs. 3C and 5A right). This observation is reasonable if we consider the working mechanism of PenG, in which the transpeptidase activity is inhibited [7, 8]. In the process involving lysozyme, such turgor was not observed. This difference can be explained by the disruption mechanisms (Fig. 5B)[5, 6, 40]. As PenG inhibits only *de novo* crosslinking of peptidoglycan, the matured parts of the peptidoglycan layer apply turgor to the cell. However, the damage by lysozyme disrupts large parts of the cell envelope.

### Change in surface pattern after *murE* operon repression

When the supply of muropeptide was stopped, the cell shape was disturbed around the cell pole (Figs. 4 and 5A right). At that time, the features of the pattern on the surface were also lost. Since the peptidoglycan layer of *B. subtilis* is 20–40 nm thick, insertion of newly synthesized filaments into the peptidoglycan layer through muropeptide supply is thought to occur on the side close to the membrane [41, 42]. If it is true, the influence on the surface pattern of the muropeptide depletion should occur in the final stage of morphological change. In fact, a noticeable change in pattern was observed when the cells were still maintaining the rod shape (Fig. 4C). This may suggest that there is fluidity in the existing peptidoglycan filaments [42, 43].

## CONCLUDING REMARKS

In this study, we showed that quick-freeze deep-etch EM is useful to visualize the peptidoglycan layer and its disruption process of *B. subtilis* in high spatial and time resolutions. Information on the structure of peptidoglycan is critical to understanding the survival strategy and evolution of bacteria in general. The application of quick-freeze deep-etch EM to peptidoglycan study should contribute to our understanding of bacteria.

## Funding

This work was supported by a Grant-in-Aid for Scientific Research on the Innovative Area “Harmonized Supramolecular Motility Machinery and Its Diversity” (MEXT KAKENHI Grant Number 24117002), by Grant-in-Aids for Scientific Research (B) and (A) (MEXT KAKENHI Grant Numbers 24390107 and 17H01544) and by the Osaka City University (OCU) Strategic Research Grant 2019 for top priority researches to MM.

## Acknowledgements

We appreciate the helpful input from Eisaku Katayama at Osaka City University and Daisuke Shiomi at Rikkyo University, gift of *B. subtilis* strains from Yoshikazu Kawai at Newcastle University, and technical support from Junko Shiomi at Osaka City University. The application of the quick-freeze, deep-etch EM technique to microbiology was developed as the general supporting project for the Grant-in-Aid for Scientific Research on Innovative Areas “Harmonized Supramolecular Motility Machinery and Its Diversity” (25117501) directed by MM.

## Author contributions

All authors contributed to the research design, analyzing data and writing the paper; I.T. and Y.O.T. performed the experiments. The authors declare no conflict of interest.

## LEGENDS

**Movie S1.** Tomogram of isolated cell for 96 degrees. The cylindrical and cell pole parts are colored differently in a still image at zero angle.

**Fig. S1.**
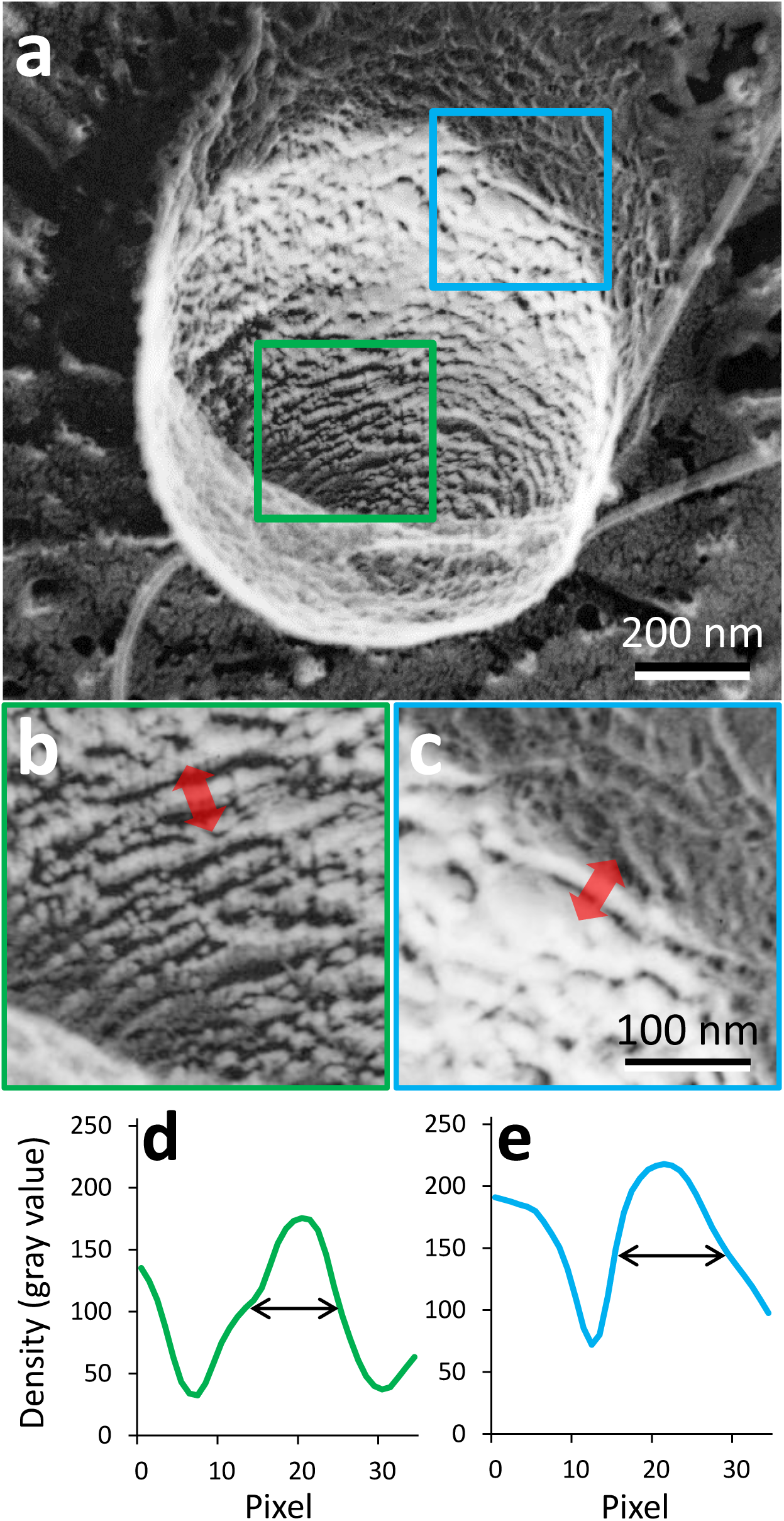
Measurement of filament width. a) Cell surface image by quick-freeze, deep-etch EM. b, c) Magnified image of areas marked by colored box in panel (a). d, e) Image profiles of red arrow area in panels (b) and (c). The positions at the mid density was defined as the boundary of focusing filament.

## Notes

#### Summary of Updates

New data and statements have been added.

